# Cleavage of the Hippo kinases and programmed cell death in murine macrophages exposed to sterile stimuli and bacterial pathogens

**DOI:** 10.1101/2025.06.15.659776

**Authors:** Sydney M. Quagliato, Ryan Mirhosiny, Lucas Meints, Matthew Gulker, Brendyn M. St. Louis, Yu-Ting Su, Pei-Chung Lee

**Author notes:** Corresponding author: Pei-Chung Lee. These authors contributed equally to this study (Ryan Mirhosiny, Lucas Meints, Matthew Gulker, Brendyn M. St. Louis).

## Abstract

Mammalian STE20-like kinases MST1 and MST2 are the conserved Hippo kinases known for their importance in organ development and tumor suppression. Notably, humans and mice lacking these kinases have increased susceptibility to infection, indicating a role of MST1/2 in immunity. In macrophages that play a critical role in host immunity, MST1/2 are proteolytically cleaved to coordinate different forms of programmed cell death, including apoptosis and pyroptosis. This cleavage event occurs when the innate immune sensors, inflammasomes, are activated by the bacterial pathogen, *Legionella pneumophila,* or damage-associated molecular patterns. In this report, we determine MST1/2 cleavage in macrophages under various inflammatory conditions and challenges with pathogenic bacteria. The sterile molecules ATP and nigericin induce MST1/2 cleavage and apoptosis when the NLRP3 inflammasome and GSDMD-mediated pyroptosis are activated. Remarkably, in conditions without NLRP3 or GSDMD activation, MST1/2 are still cleaved by caspases to promote cell death in macrophages treated with these sterile molecules. During infection, wildtype macrophages trigger MST1/2 cleavage and apoptosis against *L. pneumophila* and *Yersinia pseudotuberculosis* but preferentially activate GSDMD-mediated pyroptosis against *Pseudomonas aeruginosa* and *Salmonella enterica* Typhimurium. Interestingly, GSDMD knockout macrophages opt to cleave MST1/2 and undergo apoptosis in response to *P. aeruginosa* and *S. enterica*, suggesting an interplay between GSDMD and MST1/2. Together, macrophages funnel apoptotic death signals through MST1/2 cleavage upon stimulation of the inflammatory molecules and pathogens, which illustrates the broad implications of the host Hippo kinases in infections and sterile inflammation.

## Introduction

In the mammalian immune system, macrophages are important phagocytic cells that engulf and eliminate microbes or dead host cells. Additionally, when exposed to specific pathogen- or damage-associated molecular patterns (PAMPs/DAMPs), macrophages can activate programmed cell death pathways, including apoptosis and pyroptosis. While programmed cell death of macrophages plays a key role in pathogen restriction and inflammatory responses, different death pathways have distinct features (1–3). Pyroptosis is primarily driven by activation of gasdermin proteins by inflammasomes located within the cell cytosol. Upon sensing PAMPs or DAMPs, such as bacterial flagellin or extracellular ATP, respectively, inflammasomes activate the cysteine protease caspase-1. Consequently, caspase-1 cleaves gasdermin D (GSDMD), producing GSDMD N-terminal fragments (GSDMD-NT) which form pores in the cell membrane and result in pyroptotic cell death (4–8). In comparison, apoptosis is triggered externally by the activation of death receptors on the cell surface or internally by release of mitochondrial cytochrome c. Through a serial cleavage event by a set of pro-apoptotic caspases, including caspase-8, -9, -3, and -7, cells undergo apoptosis with cellular and molecular features, such as chromatin condensation, histone phosphorylation, cleavage of the DNA modifying enzyme poly-ADP-ribose polymerase 1 (PARP1), and formation of apoptotic bodies (9, 10). Recent studies have shown the flexibility of inflammasomes in activating the pro-apoptotic caspases and triggering PARP1 cleavage (11–20). Moreover, activated caspase-3 can contribute to lytic pyroptosis by cleaving another gasdermin protein, GSDME (21, 22).

The Hippo kinases are the core components of conserved Hippo signaling which controls cell proliferation, differentiation, and cell cycle in eukaryotes (23, 24). The <u>m</u>ammalian <u>ST</u>E20-like protein kinase-1 (MST1) and -2 (MST2) are the Hippo kinases in humans and mice and well-known for their tumor-suppressive activity (25). *Mst1/2* gene deletions lead to tumor formation in mice, and overproduction of MST1/2 enhances apoptosis of cancer cells (26–31). Importantly, anti-cancer agents induce caspase-dependent cleavage of full-length MST1/2 into smaller N-terminal fragments (MST1/2-NT) (29–33). MST1/2-NT contain the kinase domains with increased phosphorylation activity (32, 33) and become more potent in promoting apoptosis than full-length MST1/2 (30, 31). Although macrophages with MST1/2 double knockouts (DKO) are resistant to pro-apoptotic drugs (34), naïve T cells lacking MST1 easily undergo apoptosis (35–37), suggesting a complex role of MST1/2 in immune cell death. Since patients with loss-of-function mutations in *MST1* are immunocompromised (36, 37), it is necessary to understand how MST1/2 control death of different immune cell types and the inflammatory stimuli that can induce MST1/2 cleavage.

The importance of MST1/2-NT production in promoting apoptosis in macrophages with activated inflammasomes has been recently uncovered (38). Upon sensing flagellin from the bacterial pathogen *Legionella pneumophila*, the NLRC4 inflammasome activates caspase-1, leading to the cleavage of MST1/2 into MST1/2-NT. Similarly, when the NLRP3 inflammasome is activated by extracellular ATP or the potassium ionophore nigericin, MST1/2-NT production and apoptosis are readily induced. Interestingly, in conditions lacking NLRP3 activation, ATP treated macrophages still undergo programmed cell death (39, 40), implying a potential role of MST1/2-mediated apoptosis. In this report, we investigate MST1/2 cleavage and cell death in macrophages stimulated with sterile molecules or challenged with bacterial pathogens expressing different virulence factors. The results reveal that MST1/2 cleavage can be triggered by a variety of inflammatory stimuli in conditions with or without inflammasome activation and is a selective host response against pathogenic bacteria.

## Results

### ATP and nigericin induce cell death and MST1/2 cleavage in primed and non-primed macrophages

To activate the NLRP3 inflammasome in macrophages, pre-treatments with microbial ligands, known as priming, is required (41). In lipopolysaccharides (LPS) or the lipopeptide Pam3CSK4 primed immortalized bone marrow-derived macrophages (iBMDMs), ATP and nigericin induce the cleavage of both GSDMD and MST1/2 (Fig. 1A-B). Meanwhile, molecular signatures of apoptosis, including production of the activated caspase-3 p17 fragment, phosphorylation of serine 139 (p-S139) in histone H2AX, and PARP1 cleavage, were also triggered (Fig. 1A-B). These observations are consistent with prior studies in primary macrophages with activated inflammasomes (18–20, 38, 40). To determine whether MST1/2-NT production is a result of GSDMD-mediated pyroptosis, we used the chemical inhibitor dimethyl fumarate (DMF) that blocks GSDMD cleavage and pore formation (42). While production of GSDMD-NT was effectively inhibited, MST1/2-NT and the apoptosis signatures remained unchanged in primed macrophages treated with ATP or nigericin (Fig. 1A-B). The GSDMD inhibitor treatment also failed to prevent the release of lactate dehydrogenase (LDH), a cytoplasmic enzyme, into the culture media (Fig.1C-D), which aligns with the findings in previous reports using GSDMD knockout macrophages (5, 20, 38). Next, we examined the requirement of priming for MST1/2 cleavage in macrophages. Without the priming agents, neither ATP nor nigericin induced GSDMD cleavage, confirming that priming is necessary for activation of the NLRP3 inflammasome. Surprisingly, MST1/2-NT and apoptosis were still induced by ATP or nigericin in these macrophages without pre-stimulation of LPS or Pam3CSK4 (Fig. 1A-B). These non-primed macrophages also underwent cell lysis upon the ATP and nigericin treatment, although to a lesser extent than primed macrophages with activated NLRP3 (Fig. 1C-D). In the non-priming condition, ATP induced MST1/2 cleavage and cell lysis in a dose-dependent manner, while production of MST1/2-NT and the apoptotic signatures were detectable as early as one hour after exposing to ATP (Fig. 2A-B). Together, the results demonstrate that the sterile molecules can activate MST1/2-NT production and apoptosis bypassing GSDMD-mediated pyroptosis or NLRP3 activation in macrophages. In the following experiments, we focused on the non-priming condition to elucidate the effects of MST1/2 cleavage on this death route independent of NLRP3/GSDMD.

**Figure 1.**
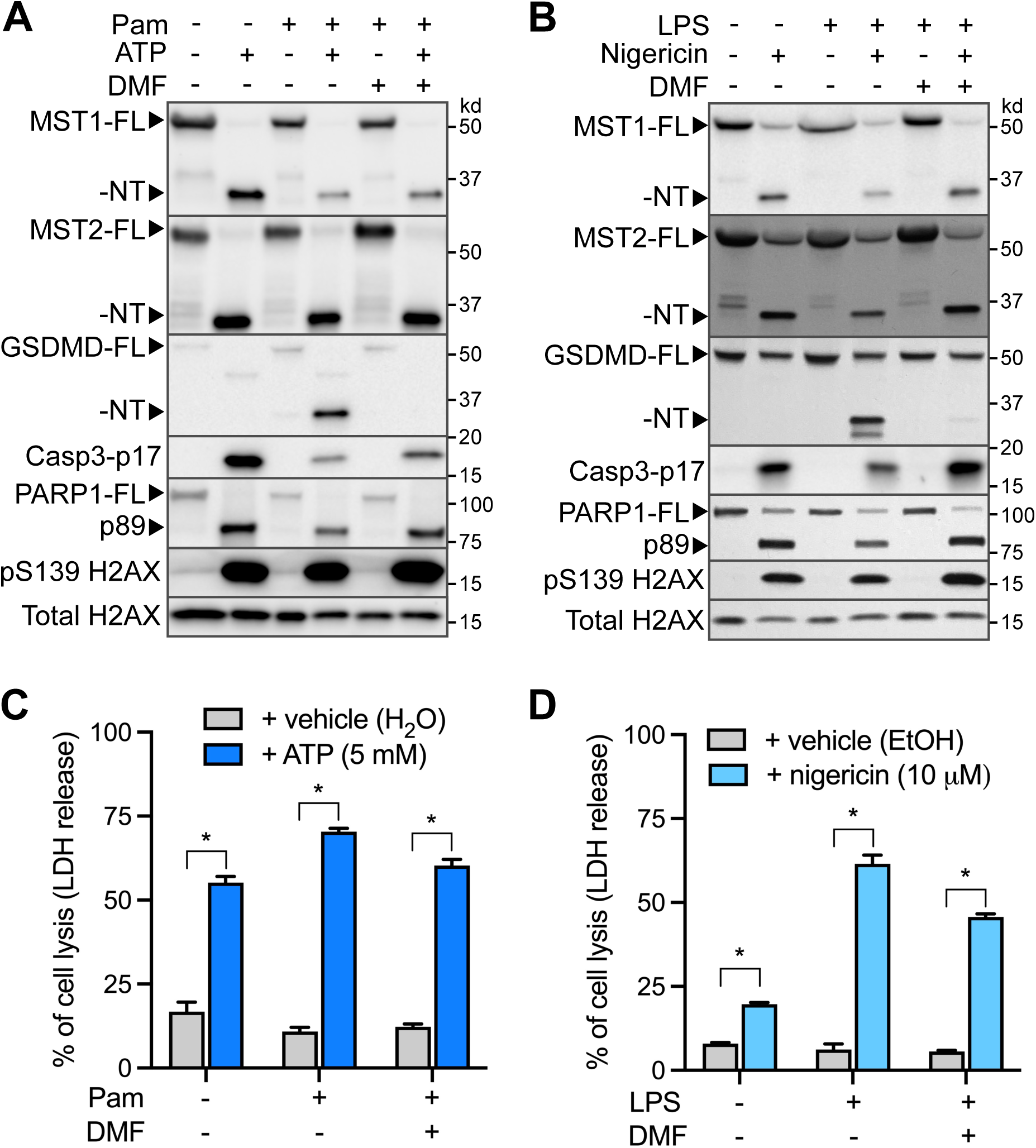
ATP and nigericin induce MST1/2 cleavage and cell death in macrophages. **A.** iBMDMs were stimulated with or without Pam3CSK4 (Pam) for 3 hours followed by the ATP (5 mM) treatment for 3 hours. DMF was added one hour before ATP stimulation. Indicated proteins in the cell lysate were determined by immunoblotting. Total Histone H2AX served as an internal control. **B.** iBMDMs were stimulated with or without LPS for 3 hours followed by the nigericin treatment (10 µM) for 4 hours. DMF was added one hour prior to nigericin. Indicated proteins in the cell lysate were determined by immunoblotting. **C.** iBMDMs were treated as in A. LDH release into conditioned media was quantified by a colorimetric cytotoxicity assay. The release levels were normalized and presented as percentages of total LDH release from iBMDMs lysed with Triton-X100. Data were representative of three independent biological repeats and presented as the mean ± SD of technical triplicates. *p<0.01, student’s t-test, two-tailed, unpaired. **D.** Cell lysis in C was determined by the LDH release assay as described in C.

**Figure 2.**
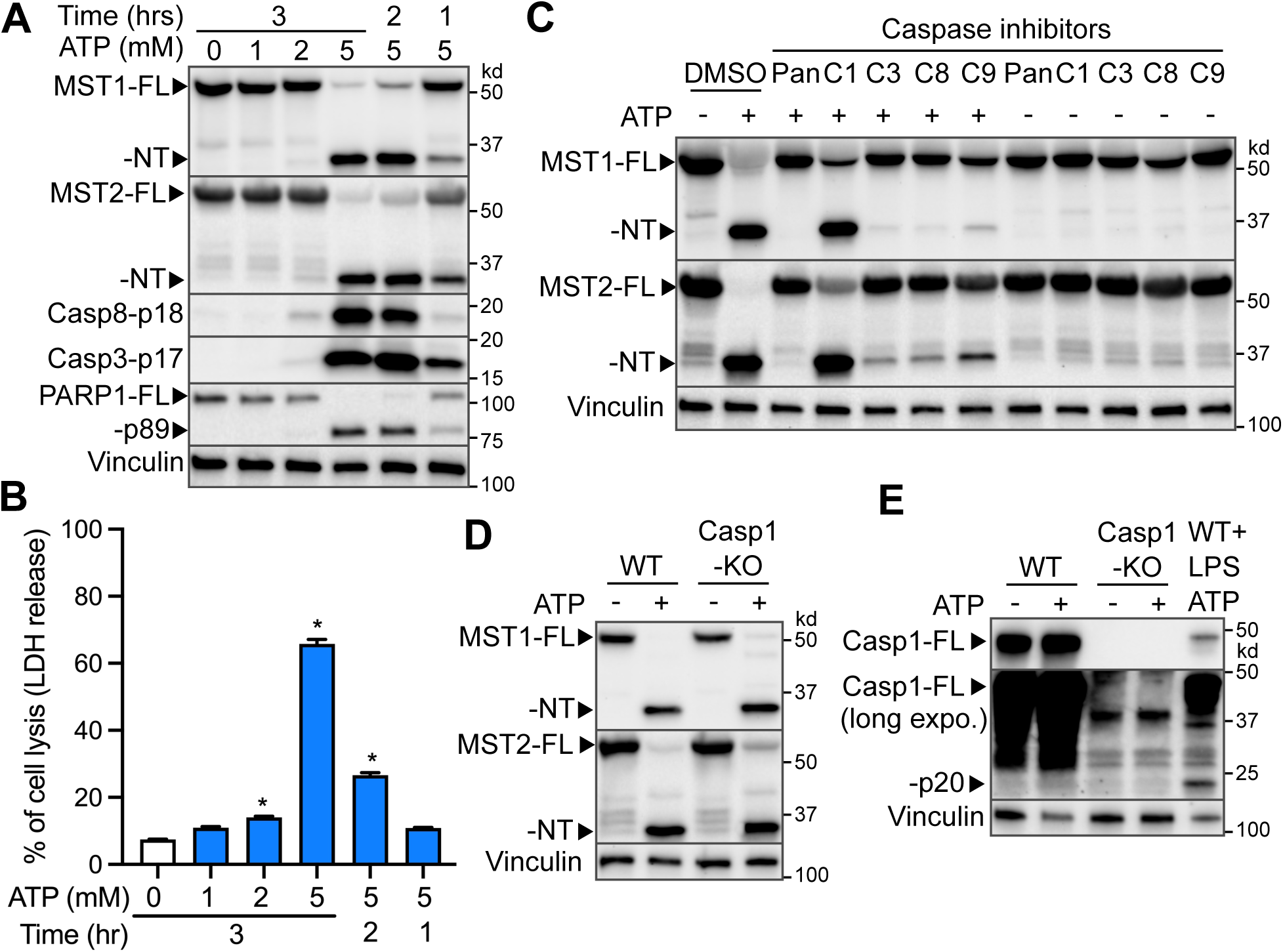
Caspases other than caspase-1 are responsible for MST1/2 cleavage in non-primed macrophages. **A.** iBMDMs were treated with ATP at indicated concentrations and time points. **B.** Cell lysis in A was determined by the LDH release assay as described in Fig. 1C. *p<0.01, compared to no ATP for 3 hours, student’s t-test, two-tailed, unpaired. **C.** iBMDMs were treated with either a caspase-1, 3, 8 or pan-caspase inhibitor 1 hour prior to ATP stimulation (5 mM) for 3 hours. **D and E.** WT or caspase-1 (Casp1) knockout iBMDMs were treated with or without ATP (5 mM) for 3 hours. WT iBMDMs primed with LPS and treated with ATP were used as a positive control of caspase-1 activation (Casp1-p20). long expo.: long exposure of the caspase-1 blot. Indicated proteins in the cell lysates were determined by immunoblotting. Vinculin served as an internal control.

### Apoptotic caspases are involved in ATP-triggered MST1/2 cleavage in non-primed macrophage

In vitro cleavage assays using recombinant proteins show that MST1 is a substrate of several caspases, including caspase-1, -3, -7, and -9 (29, 38). We sought to identify proteases that may participate in MST1/2 proteolysis in cells. Pre-treatment with the pan-caspase inhibitor Z-VAD-FMK effectively prevented ATP-induced MST1/2 cleavage in non-primed macrophages, confirming the involvement of cysteine proteases (Fig. 2C). While the levels of MST1/2 full-length (FL) proteins were partially restored, the caspase-1 specific inhibitor, VX765, had limited effects on MST1/2-NT production (Fig. 2C). To exclude the possibility that the marginal effect was due to the efficacy of VX765, we treated caspase-1 knockout (Casp1-KO) iBMDMs with ATP. Casp1-KO macrophages also had slightly increased MST1/2-FL upon the ATP treatment and produced MST1/2-NT comparable to the levels in WT macrophages (Fig. 2D), recapitulating the effects of the caspase-1 inhibitor in wildtype macrophages. Moreover, without LPS stimulation prior to the ATP treatment, active capsase-1 p20 fragment was undetectable (Fig. 2E), confirming that ATP did not activate the NLRP3 inflammasome under the non-priming condition. In contrast to VX765, the inhibitors for caspase-3, -8, and -9, although not completely, suppressed MST1/2-NT production (Fig. 2C). These findings suggest that multiple pro-apoptotic caspases are responsible for MST1/2 cleavage in non-primed macrophages treated with ATP.

### MST1/2 knockout macrophages are resistant to ATP- or nigericin-induced cell death

Our previous study demonstrates that MST1/2-NT production plays a critical role in promoting apoptosis in macrophages with activated inflammasomes (38). Since MST1/2-NT and apoptosis are triggered by ATP or nigericin without inflammasome activation (Fig. 1), we evaluated whether MST1/2 double-knockout (DKO) macrophages still undergo apoptosis. While wildtype macrophages exhibited robust MST1/2 cleavage and apoptosis in response to ATP or nigericin, the apoptotic signatures were minimal in multiple MST1/2-DKO clones under the non-primed condition (Fig. 3A-B). The DKO macrophages were also more resistant to cell lysis induced by ATP or nigericin (Fig. 3C-D), which was likely due to reduced cleavage of GSDME by caspase-3 (Fig. 3A-B). Intriguingly, while the caspase-3 inhibitor reduced MST1/2-NT production in WT macrophages (Fig. 2A), MST1/2-DKO macrophages had much lower levels of activated caspase-3 p17 when compared to WT macrophages (Fig. 3A-B). These results suggest that MST1/2-NT production by these caspases further enhances caspase-3 activation and amplify apoptosis.

**Figure 3.**
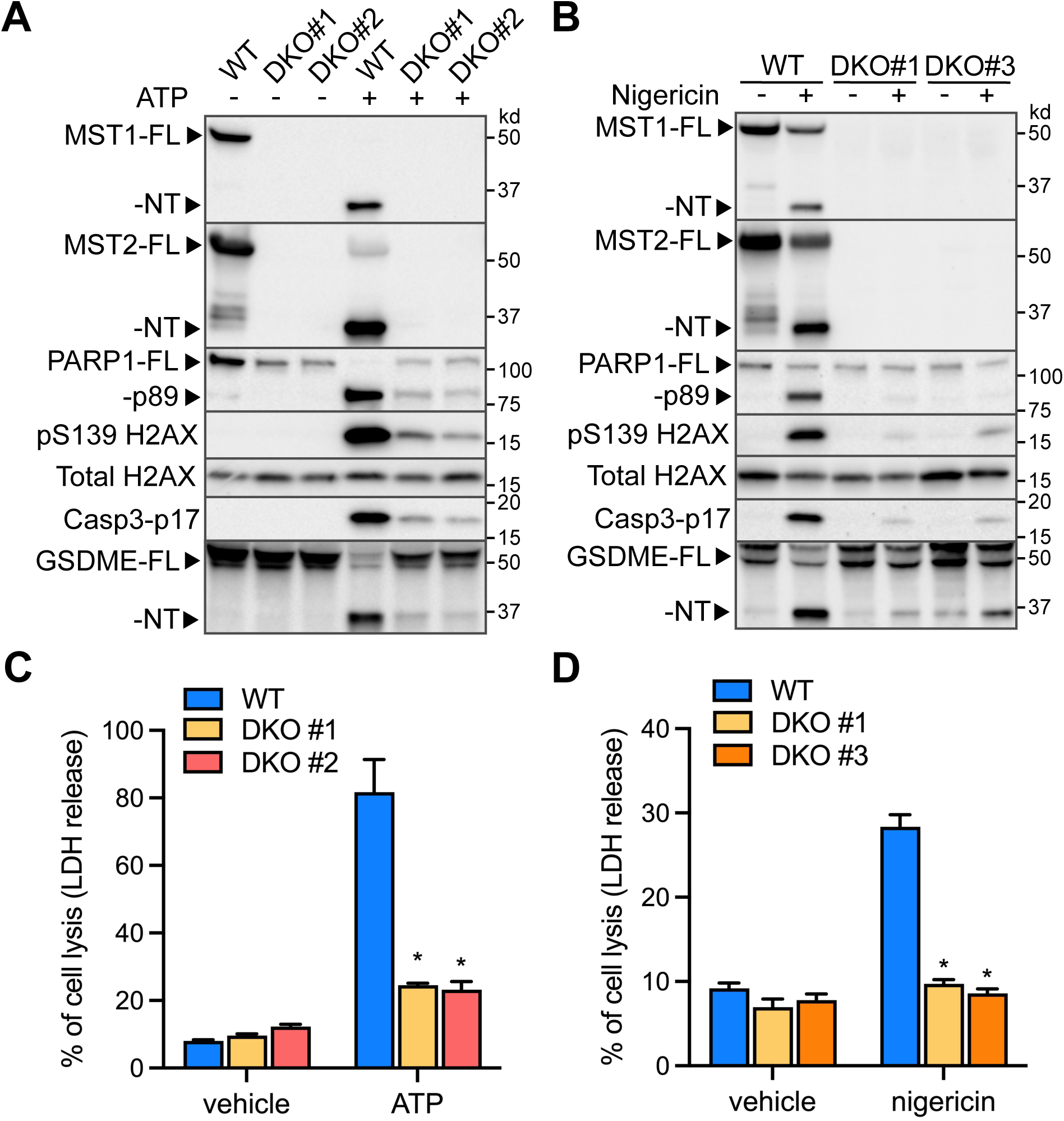
MST1/2 are required for ATP triggered macrophage cell death. **A and B.** Non-primed WT or MST1/2 double-knockout (DKO) iBMDMs (clones #1, 2, or 3) were stimulated with ATP (5 mM) for 3 hours or nigericin (10 µM) for 4 hours. Indicated proteins in the cell lysates were determined by immunoblotting. Total H2AX served as an internal control. **C and D.** LDH release of iBMDMs in A and B was determined as described in Fig. 1C. *p<0.01, compared to WT iBMDMs treated with ATP or nigericin, student’s t-test, two-tailed, unpaired.

### MST1 and MST2 have overlapped activities in promoting cell death

MST1/2 in humans and mice share high degrees of sequence identity, particularly in their kinase domains (Fig. S1). To assess their individual effects on ATP-induced cell death, we used iBMDMs that are expressing MST1 only (MST1+; MST2 knockout) or expressing MST2 only (MST2+; MST1 knockout). The single knockout macrophages triggered cell lysis in response to ATP at the levels that were significantly higher than MST1/2-DKO macrophages (Fig. 4A). In line with the increased cell lysis, MST1+ or MST2+ macrophages produced MST1-NT or MST2-NT, respectively, and the apoptotic signatures, such as caspase-3 p17 (Fig. 4B). While MST1+ macrophages fully converted MST1-FL into MST1-NT, MST2+ macrophages appeared to be less efficient in producing MST2-NT, resulting in moderate PARP1 cleavage, histone phosphorylation, and caspase-3 activation (Fig. 4B). In addition, the levels of activated caspase-3 in these macrophages were associated with the levels of GSDME-NT and cell lysis (Fig. 4A-B), indicating that activation of the pore-forming GSDME by caspase-3 may contribute to the difference in cell lysis triggered by ATP in the non-priming condition.

**Figure 4.**
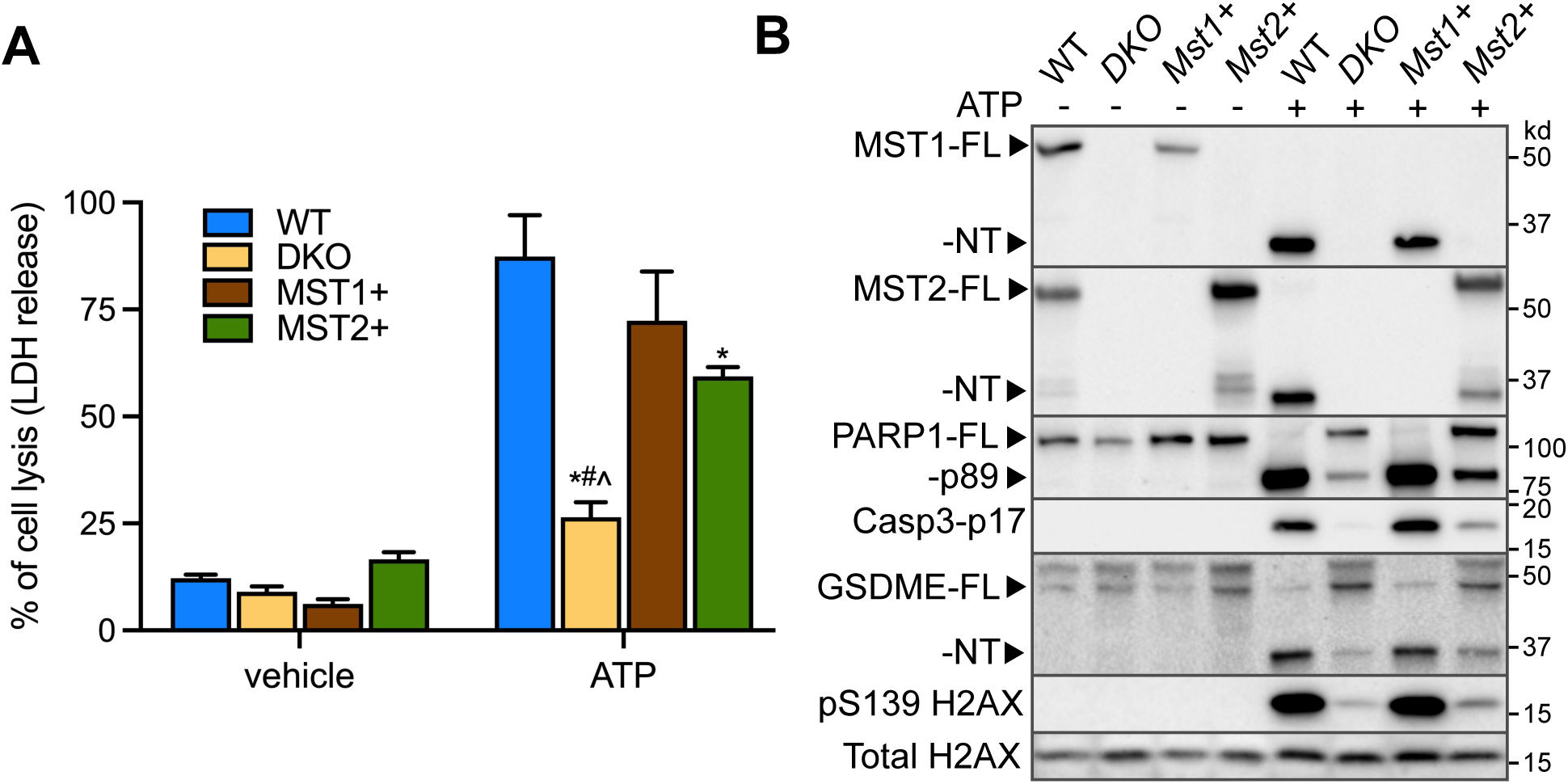
MST1 or MST2 alone is sufficient to trigger cell death in ATP-treated macrophages. **A.** WT, MST1/2 DKO, MST1-expressing (*Mst1+*, MST2 KO), MST2-expressing (*Mst2+*, MST1 KO) iBMDMs were treated with ATP (5 mM) for 3 hours and the LDH release was quantified as in Fig. 1C. p<0.01 compared to WT (*), MST1+ (#), MST2+ (^) with ATP, student’s t-test, unpaired, two-tailed. **B** WT, MST1/2 DKO*, Mst1+, Mst2+* iBMDMs were treated as in A, and indicated proteins in the cell lysates were determined by immunoblotting. Total histone H2AX served as an internal control.

### Macrophages trigger MST1/2 cleavage in response to bacterial pathogens

Since apoptosis is a critical host response during bacterial infection, we examined the status of MST1/2 in macrophages challenged with different pathogenic bacteria. As previously reported (38), *L. pneumophila* induced MST1/2-NT production and the apoptotic signatures, including PARP1 cleavage, histone phosphorylation, and caspase-8/-3 activation accompanied by GSDME cleavage. Despite the reduction in full-length MST1/2, *Pseudomonas aeruginosa* did not trigger MST1/2 cleavage or apoptosis (Fig. 5A). Likewise, MST1/2 cleavage and the apoptotic signatures were not detected in iBMDMs challenged with *Salmonella enterica* Typhimurium. RAW264.7 macrophages responded to these pathogens in similar patterns with pronounced MST1/2 cleavage and apoptosis against *L. pneumophila* while favoring GSDMD cleavage against *P. aeruginosa* or *S. enterica* (Fig. 5B). Importantly, macrophages challenged with *Yersinia pseudotuberculosis* also activated MST1/2-NT production and subsequent apoptosis (Fig. 5A), showing that MST1/2 cleavage is not limited to *L. pneumophila* infection.

**Figure 5.**
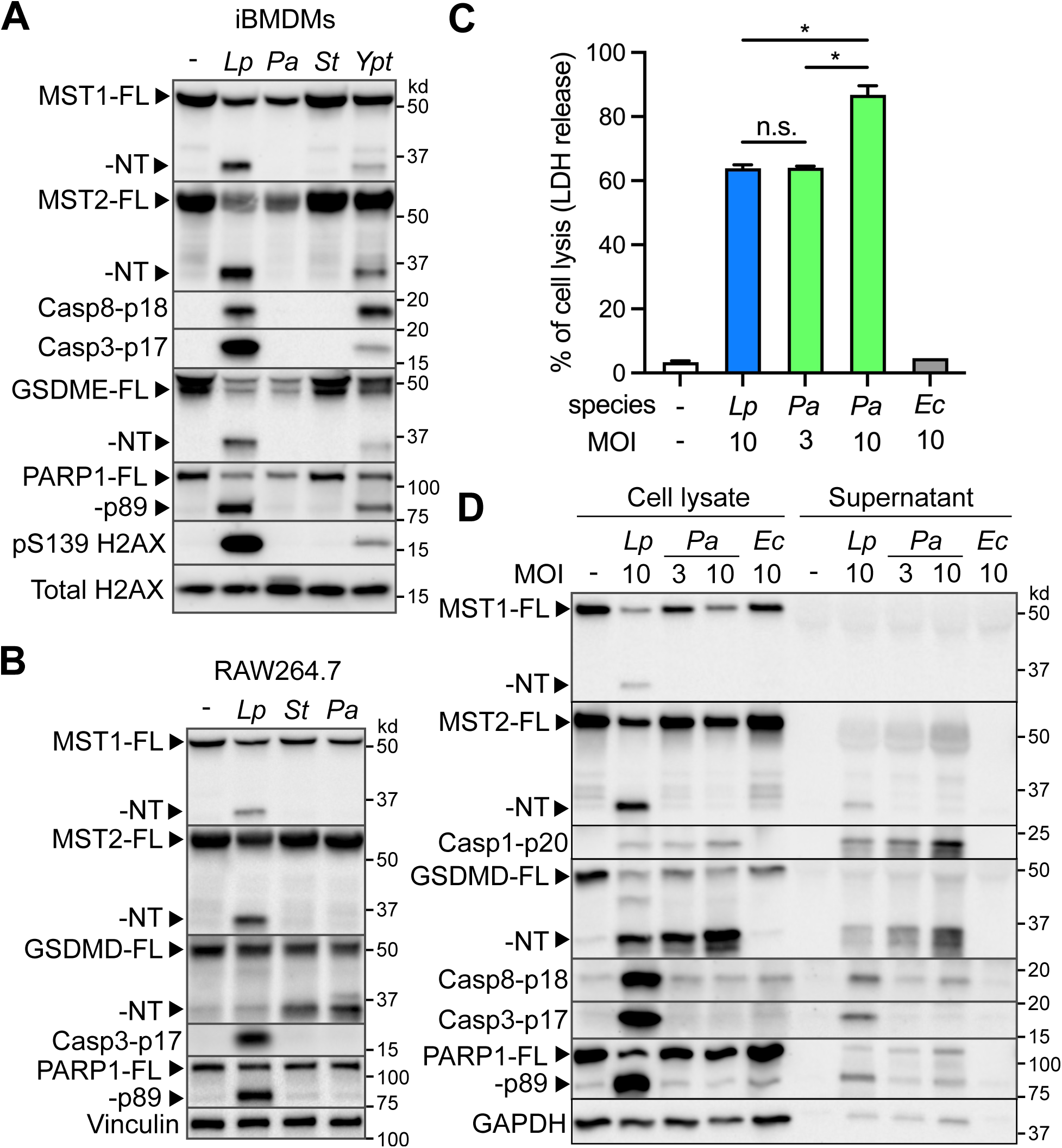
Macrophages activate MST1/2 cleavage and cell death in response to bacterial pathogens. **A.** iBMDMs were challenged with *L. pneumophila (Lp), P. aeruginosa (Pa), S. enterica* Typhimurium *(St),* or *Yersinia pseudotuberculosis (Ypt)* at MOI=10 for 3 hours. Indicated proteins in cell lysates were determined by immunoblotting. **B.** RAW264.7 macrophages were challenged with *Lp, St,* or *Pa* as described in A. **C.** iBMDMs were challenged with *Lp, Pa*, or *E. coli* (*Ec*) at indicated MOIs for 3 hours. The LDH release was quantified as described in Fig. 1C. *p<0.01, student’s t-test, unpaired, two-tailed. **D.** The cell lysate and precipitated culture supernatant samples collected from the experiment in C were analyzed by immunoblotting. Total H2AX, vinculin, GAPDH served as internal controls.

Although the same MOIs (multiplicity of infection) were used across the infection experiments to ensure similar bacterial loads, variabilities among the species may account for the observed difference in MST1/2 cleavage. To address this, we simultaneously challenged wildtype iBMDMs with *L. pneumophila* and *P. aeruginosa* at different MOIs. The LDH release assay showed that, at MOI=10, *L. pneumophila* caused lower cell lysis than *P. aeruginosa* (Fig. 5C) but was at a level comparable to *P. aeruginosa* at MOI = 3. We further analyzed the culture supernatant samples containing proteins leaked from dead cells. Since the NLRC4 inflammasome detects the flagellin proteins from *L. pneumophila* and *P. aeruginosa* and activates cell death (43–47), activated caspase-1 p20 fragments and cleaved GSDMD-NT were detected in the cell lysate and supernatant fractions upon challenge of the bacteria (Fig. 5D). Notably, while the caspase-1 p20 fragments indicated that the NLRC4 inflammasome was activated to a similar extent between *L. pneumophila* (MOI = 10) and *P. aeruginosa* (MOI = 3), MST1/2-NT and the apoptotic signatures were mainly produced in macrophages challenged with *L. pneumophila* (Fig. 5D). Moreover, non-pathogenic *Escherichia coli* did not cause cell death, caspase activation, or MST1/2 cleavage (Fig. 5D). These observations suggest a mechanism by which caspase-1 activated by the NLRC4 inflammasome can distinguish between invading bacterial species and selectively choose MST1/2 or GSDMD for cleavage.

Both MST1/2 and GSDMD are substrates of caspase-1 (5, 6, 38). We then tested whether the removal of GSDMD affects MST1/2 cleavage in macrophages. In stark contrast to wildtype iBMDMs, GSDMD-KO iBMDMs challenged with *P. aeruginosa* or *S. enterica* activated MST1/2-NT production and apoptosis (Fig. 6A). These responses were consistently triggered in both WT and GSDMD-KO macrophages challenged with *L. pneumophila*. Conversely, *L. pneumophila* induced more GSDMD cleavage in MST1/2-DKO macrophages than in WT macrophages (Fig. 6B) while the caspase-1 activity remained comparable (Fig. S2). Taken together, these results reveal an inverse relationship between MST1/2 and GSDMD cleavage, enabling the host cells to activate different death routes according to the pathogens.

**Figure 6.**
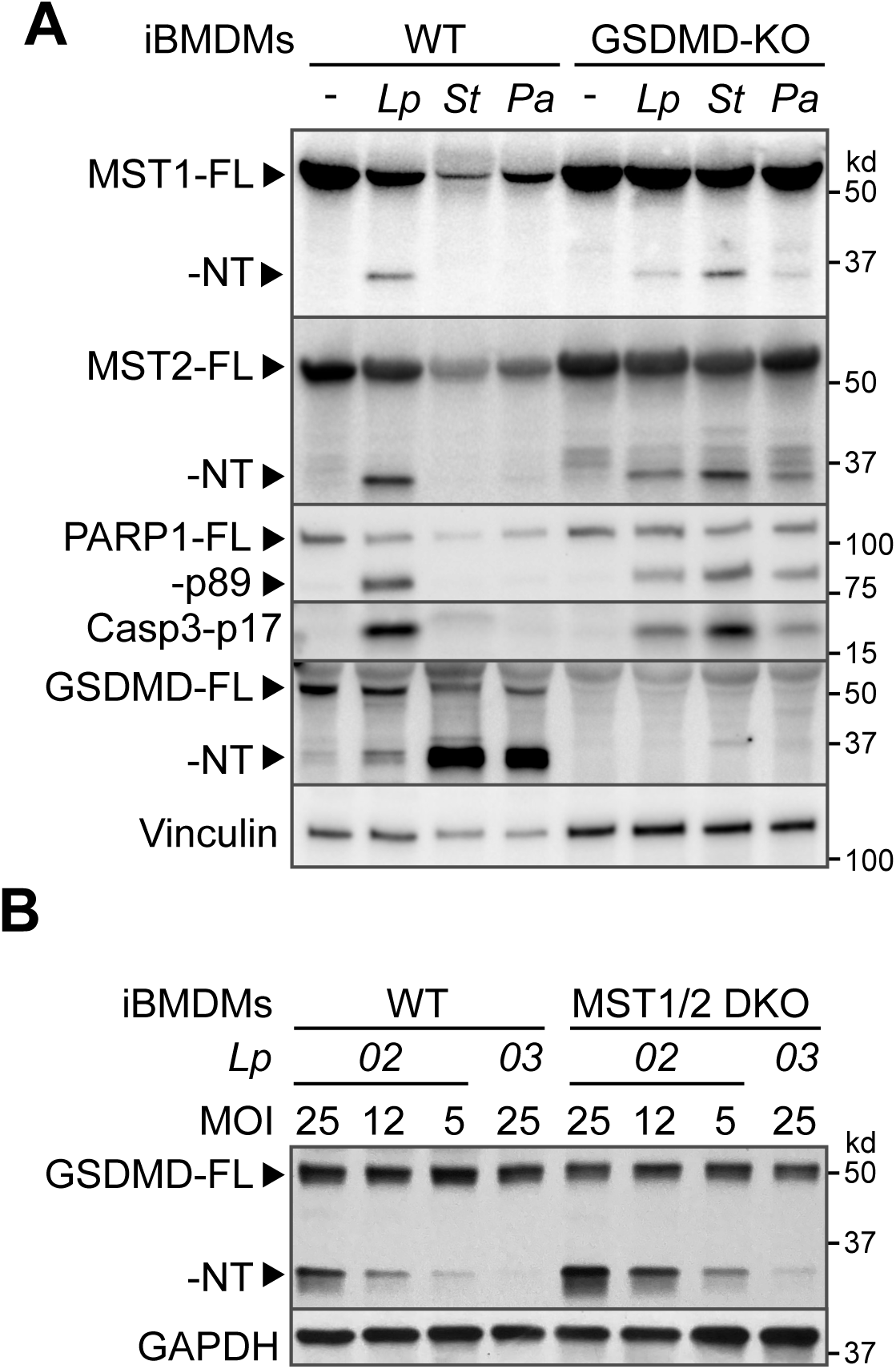
MST1/2 and GSDMD mutually influence the cleavage process. **A.** WT or GSDMD knockout iBMDMs were challenged with *Lp, Pa,* or *St* at MOI = 10 for 3 hours. **B.** WT or MST1/2 DKO iBMDMs were infected with *Lp* at MOI = 25, 12 or 5 for 3 hours. Indicated protein levels in the cell lysates were determined by immunoblotting. Vinculin and GAPDH served as internal controls.

## Discussion

As the core components of the Hippo pathway, MST1/2 are best known for influencing tissue development and tumor formation in human and mice. Innate immunity is the first line defense against pathogens and conserved among hosts. Through studying MST1/2 in macrophages, this study reveals that the proteolytic reprogramming of MST1/2 is an immune response to diverse stimuli, including damage-associated molecules and bacterial pathogens. Notably, extracellular ATP and nigericin trigger MST1/2 cleavage to promote apoptosis in conditions that do not activate GSDMD-mediated pyroptosis, indicating a role of this MST1/2 reprogramming and subsequence apoptosis in sterile inflammation.

Conversion of full-length MST1 into MST1-NT is initially characterized in cancer cells treated with pro-apoptotic agents, for example cytotrienin A and Fas ligands (32, 33). The cleavage process removes the inhibitory domain and nuclear export signal in MST1 C-terminus. Consequently, MST1-NT exhibits enhanced kinase activity and is concentrated in the nucleus where it phosphorylates S139 and S14 of histone H2AX and H2B, respectively, the hallmarks of apoptosis (48–50). In a similar manner, both MST1 and MST2 are cleaved in macrophages treated with staurosporine and Raptinal that are known apoptosis inducers (34). In addition to these chemicals, a recent study shows that macrophages cleave MST1/2 to promote apoptosis upon activation of the inflammasomes by *L. pneumophila* and the damage-associated molecule, ATP (38), suggesting the involvement of this reprogramming process in immune responses.

To activate the NLRP3 inflammasome, pre-exposure to the microbial ligands, such as LPS or the lipopeptide Pam3CSK4, is required (41). In LPS-treated macrophages, the NLRP3 inflammasome is essential for ATP-induced apoptosis (20). Intriguingly, in the absence of LPS or Pam3CSK4 stimulation, ATP and nigericin still trigger MST1/2-NT production and cell death (Fig. 1A-B), suggesting an NLRP3-independent path for MST1/2 cleavage and apoptosis. Indeed, treatment with a caspase-1 inhibitor or knocking out caspase-1 marginally affects ATP-induced MST1/2 cleavage under conditions without LPS or Pam3CSK4, as the non-priming conditions do not support activation of NLRP3 or caspase-1 (Fig. 2C-E). Together with the prior reports, these findings indicate that MST1/2 are integrated in two separate death routes in response to ATP and nigericin. In sterile environments lacking microbial ligands, macrophages primarily use the NLRP3-independent path to cleave MST1/2 for triggering apoptosis that is limited inflammatory. When microbial ligands are present, macrophages switch to the NLRP3/caspase-1 cascade to activate not only MST1/2-mediated apoptosis but also GSDMD-mediated pyroptosis (20, 38).

In agreement with this idea, activated caspase-3 and caspase-8 are detected in non-primed macrophages treated with ATP (Fig. 2A), and MST1/2-NT production is largely blocked by the specific inhibitors for caspase-3, -8, and -9 (Fig. 2C). The data show that these apoptotic caspases participate in MST1/2 cleavage and are consistent with the report by Zeng et al., which describes activation of the pro-apoptotic caspases by ATP in primary mouse macrophages and RAW264.7 cells without activated NLRP3 inflammasomes (40). ATP-induced cell lysis under the non-priming condition (Fig. 1A, 2B) also recapitulates the phenomenon in peritoneal macrophages reported by Le Feuvre et al., (39). This study further demonstrates that these treatments trigger MST1/2 cleavage through the NLRP3-independent path to promote apoptosis. Notably, the apoptotic signatures, including activated caspase-3 p17, are minimal in MST1/2 DKO macrophages treated with ATP or nigericin (Fig. 3A-B). The low level of capase-3 activation leads to decreased GSDME cleavage, a pore-forming gasdermin protein and known substrate of caspase-3. This likely contributes to the reduction of cell lysis in MST1/2-DKO macrophages treated with ATP or nigericin (Fig. 3C-D), indicating that MST1/2 may influence lytic death by controlling the caspase-3/GSDME cascade in macrophages. Additionally, the results in Fig. 2 and Fig. 3 reveal an intriguing mechanism by which MST1/2 cleavage and the apoptotic caspases may form a positive feedback loop to amplify apoptosis in macrophages.

Distinct from non-vertebrates and single cell eukaryotes, mammals encode two Hippo kinases (51). In MST1 and MST2 single knockout macrophages treated with ATP, the apoptotic signatures and cell lysis are restored (Fig. 4), suggesting that the two kinases have redundant functions. Upon ATP stimulation, MST1+ macrophages process MST1 cleavage efficiently and undergo cell death at similar levels as WT macrophages expressing both MST1 and MST2. It is noteworthy that MST2+ macrophages trigger intermediate levels of the apoptotic signatures and cell lysis, while noticeable MST2 full-length proteins are present and less MST2-NT is produced (Fig. 4B). The reduced cell death in MST1/2-DKO macrophages is unlikely caused by lack of the ATP-sensing receptor P2X7 receptor considering that the RNA transcripts of this gene are comparable among wildtype and MST1/2-DKO macrophages (Fig. S3). In addition, MST1/2-DKO macrophages are also resistant to apoptosis induced by nigericin (Fig. 3B, 3D) which is a potassium ionophore and does not bind to the P2X7 receptor. Together, the strong association between the levels of MST1/2-NT production and apoptosis in ATP-treated macrophages reflects the fact that preventing MST1/2 cleavage reduces apoptosis (38).

Macrophages are professional phagocytes that play a crucial role in host defense against pathogens and may undergo different forms of programmed cell death during infection. In this study, we further characterized MST1/2 cleavage in macrophages challenged with different bacterial pathogens. *L. pneumophila* induces robust MST1/2 cleavage and apoptosis along with moderate GSDMD cleavage in both iBMDMs and RAW264.7 macrophages (Fig. 5A-B), which has been attributed to the detection of *L. pneumophila* by the NLRC4 inflammasome and subsequent activation of caspase-1 (38). Strikingly, *P. aeruginosa* triggers high levels of GSDMD-NT instead of MST1/2 cleavage or apoptosis, although caspase-1 is equally activated by both pathogens (Fig. 5D). To investigate this discrepancy, we tested if it could be due to the infection styles given that *L. pneumophila* is an intracellular pathogen and *P. aeruginosa* primarily remains extracellular during infection. Intriguingly, macrophages challenged with the intracellular bacterium *S. enterica* Typhimurium preferentially cleave GSDMD over MST1/2 (Fig. 5A-B), indicating that the intracellular infection style might not be the contributing factor.

Another possibility is that macrophages distinguish between the types of protein secretion system in these pathogens. To inject effectors into host cells, *P. aeruginosa* and *S. enterica* use the type III secretion systems, whereas *L. pneumophila* uses the dot/icm type IV secretion system. Remarkably, *Y. pseudotuberculosis*, an extracellular pathogen encoding a type III secretion system induces MST1/2-NT production and apoptosis (Fig. 5A), while non-pathogenic *E. coil* does not trigger cleavage of either MST1/2 or GSDMD (Fig. 5D). Since the *Y. pseudotuberculosis* effectors, YopM and YopK, prevent caspase-1 activation by the inflammasomes (52–54), MST1/2 cleavage in response to this bacterium is likely triggered by a host immune pathway independent of the inflammasomes. Collectively, these infection experiments reveal that MST1/2 cleavage is a selective host response for promoting apoptosis in macrophages based on the species of invading pathogens.

Although the determinant(s), whether from the host cells or pathogens, for selection of the cleavage substrates requires further investigation, MST1/2-NT production and apoptosis occur in GSDMD-KO macrophages challenged with *P. aeruginosa* and *S. enterica* (Fig. 6A). Reciprocally, GSDMD cleavage is increased in MST1/2-DKO macrophages challenged with *L. pneumophila* (Fig. 6B). These results suggest a dynamic interplay between MST1/2 and GSDMD that enables macrophages to cleave the alternative substrate to activate programmed cell death when the preferred substrate is absent or targeted by the pathogen. In summary, this study demonstrates that sterile inflammatory molecules, such as ATP released from damaged cells, and bacterial pathogens are triggers for MST1/2 cleavage and apoptosis in macrophages. Notably, dysregulated MST1/2 cleavage has been reported in several human diseases. For example, liver tumor cells do not produce MST1/2-NT, whereas pancreatic β-islet cells exposed to pro-diabetic agents cleave MST1 and undergo apoptosis (26, 55). Thus, identifying the sterile stimuli and pathogens for triggering MST1/2 cleavage lays a foundation for future research in the role of the conversed Hippo kinases in sterile inflammatory disorders and infectious diseases.

## Experimental Procedures

### Mammalian cell culture

Wildtype, MST1/2 double knockout, MST2 knockout *(Mst1+*), MST1 knockout (*Mst2+*), caspase-1 knockout, and GSDMD knockout immortalized bone marrow derived macrophages (iBMDMs) (38) were cultured in DMEM (Corning, 15013CV) supplemented with 10% fetal bovine serum, 2 mM L-glutamine (Corning, MT25005CI), and penicillin/streptomycin (Corning, MT3000CI) at 37°C in humidified incubators with 5% CO_2_. RAW264.7 cells were cultured in the same conditions as iBMDMs.

### Macrophages Priming and the ATP/nigericin Treatments

iBMDMs were seeded in 12-well or 24-well plates and cultured overnight at 37°C in humidified incubators with 5% CO_2_. For priming macrophages, the original medium was removed and replaced with complete DMEM medium containing either 2 µg/mL of lipopolysaccharide (Invivogen, tlrl-eblps) or 1 µg/mL of Pam3CSK4 (Invivogen, tlrl-pms). After 3 hours of priming, ATP (final concentration: 5 mM, Millipore Sigma, A2383) or nigericin (final concentration: 10 µM, Invivogen, tlrl-nig) was added, and the cells were incubated with ATP for 3 hours or with nigericin for 4 hours. In the GSDMD inhibitor experiments, 25 µM of Dimethyl fumarate (DMF; Thermo Scientific 222180250) was added 1 hour prior to the ATP or nigericin treatments. Following treatments, cell supernatant was collected and centrifuged at 200x g for 5 minutes, while 1x Laemmli sample buffer was added to lyse attached cell monolayers in the wells. Spun media supernatant was then used for the LDH release assays. The lysed cell monolayers in 1x Laemmli sample buffer were then combined with the corresponding cell pellet in the centrifuged tubes. Combined protein samples were then denatured at 97°C for 10 minutes and analyzed by immunoblotting. For the caspase inhibitor treatments, wildtype iBMDMs were treated with 5 mM ATP for 3 hours without priming. One hour prior to ATP stimulation, cells were pre-treated with 100 µM of caspase-1 (VX765; Invivogen, inh-vx765i-1), caspase-3 (Z-DEVD-FMK; R&D Systems, FMK004), caspase-8 (Z-IETD-FMK; R&D Systems, FMK 007), or caspase-9 (Z-LEHD-FMK; R&D Systems, FMK008) inhibitor or 25 µM of pan-caspase Inhibitor (Z-VAD-FMK; R&D Systems, FMK001). Cell lysate samples for immunoblotting were collected as previously described.

### Immunoblotting

Denatured protein samples in 1x Laemmli sample buffer were separated by SDS-PAGE, transferred to a nitrocellulose membrane. Indicated proteins were detected by specific primary antibodies followed by HRP-conjugated secondary antibodies, and chemiluminescence was detected in ChemiDoc MP Imaging System (BioRad) or on X-ray films. Antibodies used in this study: Vinculin (sc-73514, Santa Cruz Biotechnology), cofilin-1 (sc-53934, Santa Cruz Biotechnology), MST2 (ab52641, Abcam), GSDMD (ab209845, Abcam), GSDME (ab215191, Abcam), MST1 (14946S, Cell Signaling), PARP1 (9542S, Cell Signaling), cleaved caspase-3 (9661S, Cell Signaling), histone H2AX (2595S, Cell Signaling), phospho-S139 H2AX (2577S, Cell Signaling), cleaved caspase-8 (8592S, Cell Signaling), caspase-1 (AG-20B-004-C100, Adipogen), goat anti-rabbit HRP-conjugated antibody (A-21244, Invitrogen), goat anti-mouse HRP-conjugated secondary antibody (G-21040, invitrogen). All immunoblots are representative of at least three independent biological repeats.

### Colorimetric LDH release assays

Release of the lactate dehydrogenase (LDH) into culture medium from control and treated macrophages were analyzed by using CytoTox 96 non-radioactive cytotoxicity assays (Promega G1780). The absorbance at 490 nm was measured using a SpectraMax i3X plate reader (Molecular Devices). All samples were analyzed in triplicate and normalized as percentages of the total cell lysate from the maximum LDH release control (iBMDMs lysed with 0.02% Triton-X100). Data from the LDH release assays are presented as mean ± SD of three technical triplicates and representative of at least three biological repeats.

### Bacterial cultures

The *L. pneumophila* (*Lp*) strain *Lp02* carrying the pJB908 plasmid was grown on CAYE agar plates, and the overnight *Lp* cultures were set up in liquid AYE media as described (56). Prior to infection, cultures were spun down at 11,000x g for 2 minutes, resuspended in DMEM (10% FBS and 2 mM L-Glutamine without penicillin or streptomycin), and diluted based on the optical density (OD_600_) of the overnight culture. The *P. aeruginosa* PAO1F strain was inoculated in high salt LB broth and cultured overnight as described (57). Overnight cultures were diluted at 1:100 and 1:50 dilutions and incubated for 2-3 hours at 37°C in a shaking incubator. The fresh cultures were prepared for infection as for *L. pneumophila*. *Salmonella enterica* serovar Typhimurium strain was received from Wayne State University’s Microbiology Lab. *S. enterica* was inoculated in regular LB broth (10 g/L tryptone, 10 g/L NaCl, 5 g/L yeast extract) and cultured overnight at 37°C in a shaking incubator. Overnight *S. enterica* cultures were diluted and prepared for infection as for *P. aeruginosa*. *Yersinia pseudotuberculosis* was a gift from the lab of Dr. Joan Mecsas, Tuft University, and cultured as described (58).

### Bacterial infection in macrophages

Macrophages were seeded in a 12-well plate one day prior to infection. One hour before infection, media was removed from the wells and replaced with 0.9 mL of DMEM (10% FBS, 2 mM L-glutamine, without penicillin or streptomycin). 100 µL of bacterial suspension was added to each well at calculated multiplicity of infection (MOIs). Plates were centrifuged at 200x g for 5 minutes, then incubated at 37°C in a humidified CO_2_ incubator. After one hour incubation, 300 µg/mL gentamicin was added into the wells, and the plates were incubated for additional 2 hours (total infection time of 3 hours). Culture media for the LDH release assay and cell lysate protein samples were collected as described for the ATP/nigericin treatment. To prepare proteins in the culture media for immunoblotting, DMEM containing 1% FBS and 2 mM L-glutamine without penicillin or streptomycin was used in the infection experiments. After infection, 450 µL of the culture supernatant was mixed with 50 µL of 100% trichloroacetic acid (TCA), and the precipitated protein pellets were washed with acetone twice. Air-dried protein pellets were resuspended in 1x Laemmli sample buffer and denatured on a heat block.

### Caspase-1 activity assays

The activity of caspase-1 in wildtype or MST1/2 double knockout iBMDMs after challenge of *L. pneumophila* at MOI = 10 for 3 hours was analyzed by the Caspase-Glo^®^ 1 Inflammasome Assay (G9951, Promega) according to the manufacturer’s manual.

### Statistical analyses

All statistical analyses and graphs were performed and produced in GraphPad Prism.

## Data availability

All data are included in this article and the supporting information.

## Supporting information

This article contains supporting information.

## Acknowledgements

The authors thank the anonymous reviewers for their constructive feedback and Dr. Yuan He (Wayne State University) for assistance with the ATP and nigericin treatments.

## Funding and additional information

This study was supported by the startup funds from Wayne State University (to P.-C.L.) and the AI186288 grant from the National Institutes of Health (to P.-C.L.). B.M.S. was supported by the Rumble Fellowship (2024–25) from the Graduate School, Wayne State University. The content is solely the responsibility of the authors and does not necessarily represent the official views of the National Institutes of Health.

