## Supplemental Figures for "Cleavage of the Hippo kinases and programmed cell death in murine macrophages exposed to sterile stimuli and bacterial pathogens"

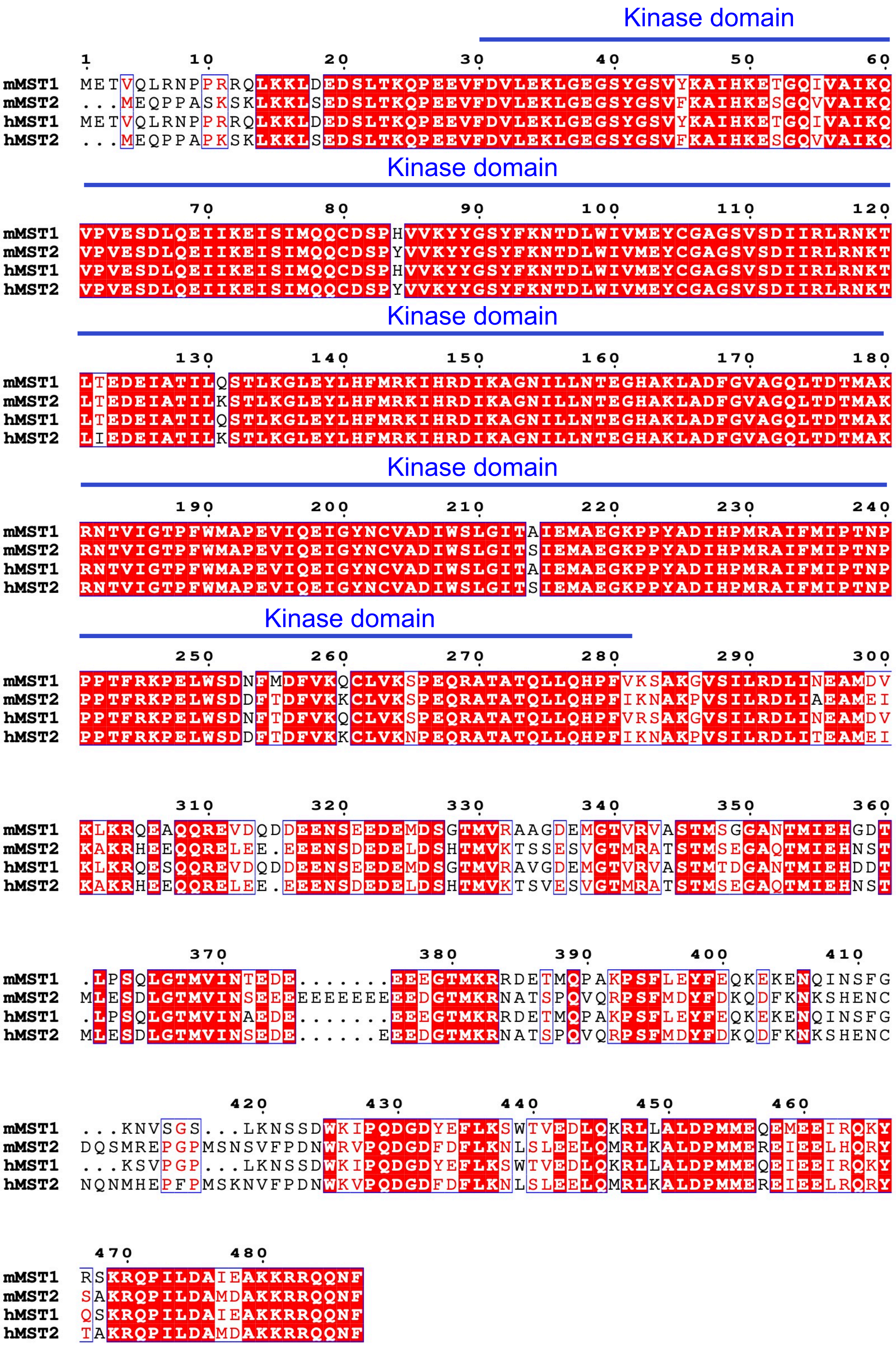


**Figure S1.** The multiple sequence alignments of mouse and human MST1/2 were processed by Clustal Omega and ESPript 3.0. Conserved amino acids were in white fonts and red boxes. Similar amino acids were in red letters and white boxes. NCBI reference sequences used: mouse MST1 (mMST1; NP_067395.1), mouse MST2 (mMST2; NP_062609.2), human MST1 (hMST1; NP_006273.1), human MST2 (hMST2; NP_006272.2).

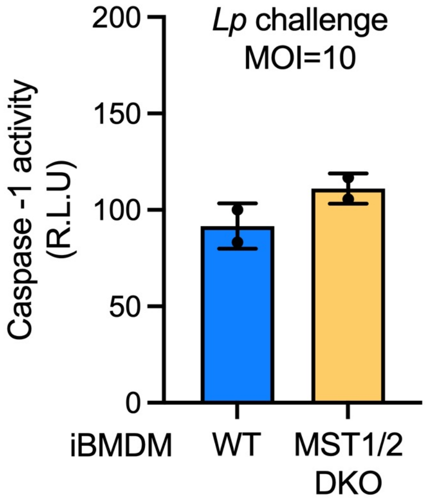


**Figure S2.** The caspase-1 activity in WT and MST1/2 DKO iBMDMs challenged with *L. pneumophila* (*Lp*) at MOI = 10 for 3 hours was measured by the luciferin-conjugated substrate activity assay (caspase-1 Glo, Promega). Data were presented as mean ± SD of two independent repeats.

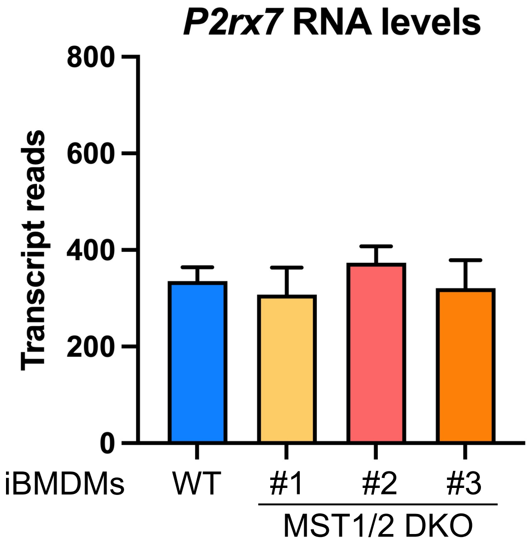


**Figure S3.** The RNA expression levels of the ATP-sensing receptor gene *P2rx7* in wildtype iBMDMs and MST1/2 DKO iBMDMs (clone #1, #2, and #3). Reads of the *P2rx7* RNA transcripts were extracted from the RNA sequencing results in *St. Louis et al., mBio 2024*, Reference (34), and presented as mean ± SD of five independent repeats. No statistical significance detected among wildtype iBMDMs and the MST1/2-DKO clones.
